# Phagosome Maturation in Macrophages is Enhanced by p38α MAPK Signaling

**DOI:** 10.1101/2025.07.03.662985

**Authors:** Mitali Shah, Nikhita Kirthivasan, Sandip Chakraborty, Yamuna Krishnan, Siddharth Jhunjhunwala

## Abstract

In macrophages, it is generally assumed that without active escape mechanisms, all phagocytosed cargo eventually reaches lysosomes. Yet, the influence of specific ligands, like lipopolysaccharide (LPS), on phagosome maturation is unclear. Here, using sterile, non-immunogenic particles as model cargo phagocytosed by macrophages, we show that in the absence of an external signal, less than half the phagosomes fuse with lysosomes. By quantifying phagosome maturation in terms of lysosomal localization and mapping lumenal pH dynamics, we find that early events triggered by LPS-induced signaling enhance both cargo delivery to lysosomes and phagosome acidification rates. We demonstrate that stress-activated p38 alpha mitogen-activated protein kinase (p38 MAPK) is important for this signaling event. Other signals known to activate p38 MAPK such as flagellin, IgG and albumin also enhance lysosomal delivery of phagocytosed cargo. Our work indicates that the maturation of phagosomes in macrophages is enhanced by the presence of specific ligands on the cargo, which activate cell surface receptors that signal via p38 MAPK.

## Introduction

Phagosome maturation is essential for the degradation of phagocytosed cargo. Phagosome maturation is characterized by the acidification of the phagosome lumen via sequential fusion events with endo-lysosomal compartments of the cell to eventually form the degradative phagolysosome (Nguyen and Yates, 2021). The dynamics of phagosome maturation are regulated and influenced by several intra- and extracellular factors, giving rise to a remarkable heterogeneity in how the process unfolds across various cell types, functional states, immune contexts and target chemistries (Pauwels et al., 2017; Jankowski et al., 2002; Fountain et al., 2021; Westman and Grinstein, 2021).

While it is acknowledged that the regulation and kinetics of phagosome maturation are context-dependent, the maturation of phagosomes into phagolysosomes is generally considered a de facto process (Fountain et al., 2021). However, the role of additional signals that influence phagosome maturation in phagocytes, such as macrophages, is unclear (Pauwels et al., 2017; Westman and Grinstein, 2021). For instance, Blander and Medzhitov suggest that in the absence of TLR-signaling, the maturation of phagosomes containing bacteria is impaired (Blander and Medzhitov, 2004), while Russell and colleagues have argued that phagosome maturation proceeds independent of TLR-signaling, but sustained signaling brings about differences in extents of acidification (Yates and Russell, 2005; Yates et al., 2007). Dill et al. follow up on these studies using a reductionist approach of particles labelled with single ligands and demonstrated that phagosome proteome changes based on ligand-cell surface receptor interaction, but they did not evaluate phagosome acidification (Dill et al., 2015). While the role of signals originating from the cell surface in phagosome maturation is debated, it is believed that all phagosomes would fuse with lysosomes eventually, but this remains to be verified.

We therefore designed a precisely controlled phagocytic system in macrophages, employing fluorescent particulates of non-biological origin, to evaluate the effects of individual cues on phagosome maturation resulting in phago-lysosomal fusion. As these particles are uniform, non-immunostimulatory, easy to track inside a cell, and amenable to surface modifications, we can precisely study the effect of a signal on phagosome maturation using automated image analysis. We find that phagosome maturation in macrophages is in fact, not de facto, and that p38 MAPK signaling is a key pathway that promotes maturation.

## Results

### Particles mature to the lysosomes in a stochastic manner after phagocytosis

We first evaluated basal levels of phagosome maturation in our model system of RAW 264.7 cells and 500 nm-sized, non-immunostimulatory, fluorescent polystyrene particles (Sharma et al., 2019, 2022). Particles were first internalized by RAW 264.7 cells over a 1-hour incubation period (pulse). Thereafter, non-internalized particles were washed off thoroughly and then cells were imaged at varying lengths of time post phagocytosis (chase) and the images were analysed for colocalisation of particles with lysosomal markers (Fig 1 A). Upon phagocytic uptake, we find that fewer than 50% particles that were internalized per cell finally colocalized with the lysosomal marker, lysosomal-associated membrane protein 1 (LAMP1) (Fig 1 B, C), or with acidified compartments that are tracked using LysoTracker Red DND-99 (LTR) (Fig 1 D, E), as analyzed using the Pearson’s correlation coefficient (PCC)-based method of analyzing colocalizstion. This observation held true across all time points tested and was agnostic of the method used – PCC or object overlap (OO)-based method – to analyze colocalization (Fig S1 A). We verified that both methods measured true colocalization robustly, by performing the same analyses on image stacks that had the lysosome channel flipped either horizontally or vertically (Fig S1 B – D). Additionally, the results from each method were recapitulated by manual analysis of colocalization (Fig S1 E – F). Henceforth, we use the OO based-method to quantify percentage colocalization of particles with lysosomes in LTR stained cells, whereas the PCC-based method is used for cells that were stained for LAMP1.

**Figure 1:**
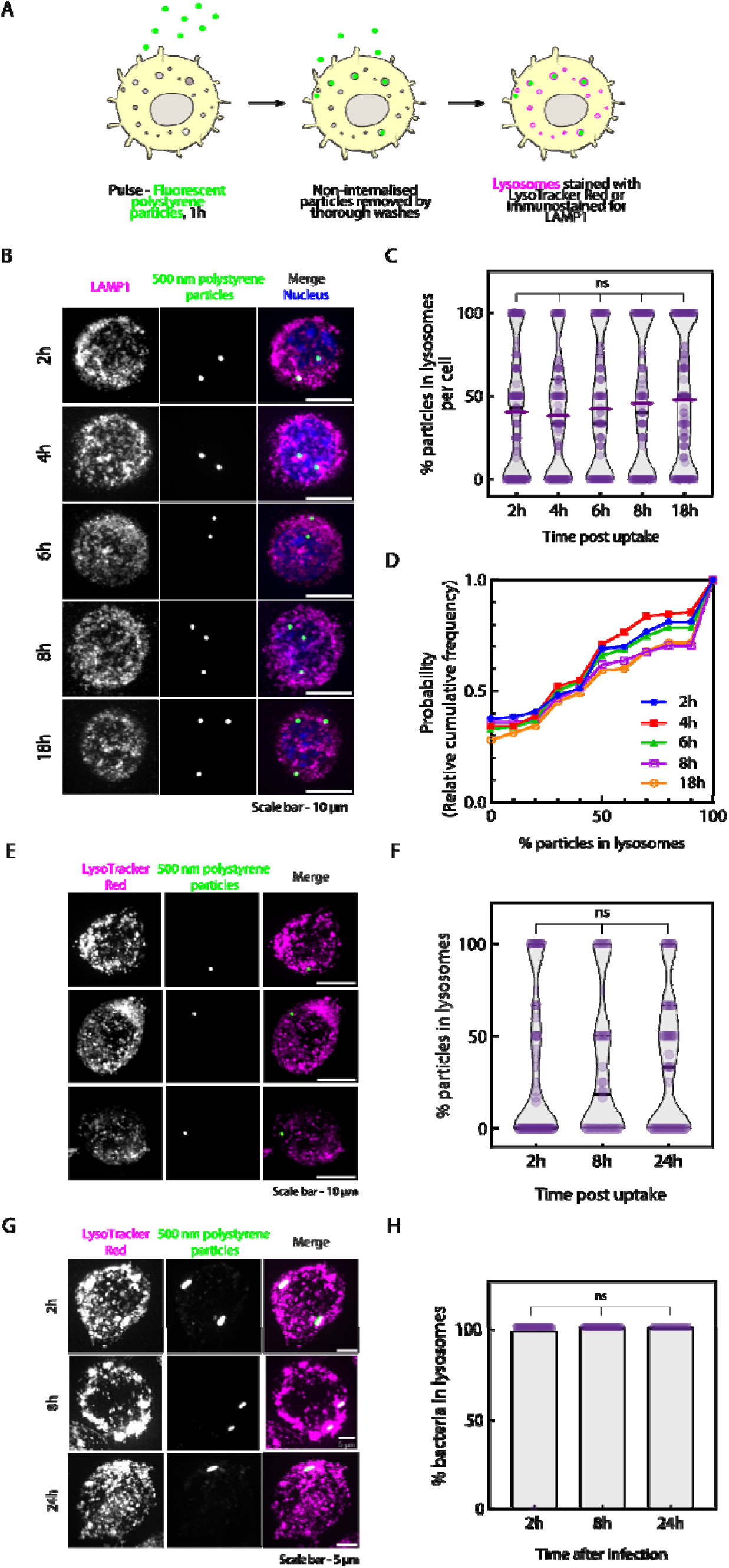
Fewer than 50% of phagocytosed particles are localized in macrophage lysosomes. **A** – Schematic describing the design of experiment prior to microscopy. **B –** Representative maximum intensity projections (MIPs) of RAW 264.7 cells at indicated time points after introduction of particles. Magenta – LAMP1, Green – 500 nm-sized non-modified polystyrene particles, Blue – Hoechst. Scale bar – 10 μm. **C –** Percentage of internalised particles per cell that colocalize with LAMP1 signal at indicated time points. Kruskal-Wallis test, N ≥ 3 independent experiments, 116 cells per group on average. Data represented as violin plots with colored lines indicating mean, black lines indicating median and dashed lines indicating quartiles of the data. **D –** Representative MIPs of RAW 264.7 cells stained with LysoTracker Red (LTR) at indicated time points after introduction of particles. Magenta – LTR, Green – 500 nm-sized non-modified polystyrene particles. Scale bar – 10 μm. **E –** Percentage of internalised particles per cell that colocalize with LTR signal at indicated time points, as obtained using the PCC-based method for analysis of colocalisation. Kruskal-Wallis test, N = 3 independent experiments, 55 cells per group on average. Data represented as violin plots with colored lines indicating mean, dark lines indicating median and dotted lines indicating quartiles of the data. **F –** Representative MIPs of RAW 264.7 cells at indicated times post infection with *E. coli* (MOI = 10:1). Magenta – LTR, Green – EGFP-expressing *E. coli.* Scale bar – 5 μm. **G –** Percentage of bacteria taken up by a cell that were localized in lysosomes. Kruskal-Wallis test, N = 3 independent experiments, n = 47 cells per group on average. Bars indicate the arithmetic mean of the data.

The percentage of internalized particles that colocalized with lysosomes did not correlate with the phagocytic capacity of the cell or the lysosomal volume (Fig S1 G, H). We also did not observe a significant difference in the number of internalized particles across cell populations at the end of different chase durations (Fig S1 I, J). Lysosomal volumes of cells which had phagocytosed particles did not differ from those that did not, except for an increase at 2 hours post uptake (Fig S1 K - data shown only for 2 hours). Even when pooling all analyzed particles across cells, we found ∼42% of all particles colocalized with lysosomes 2 hours post uptake (Fig S1L). By conjugating pHrodo to the particle surface, we determined that the average phagosomal pH was 6.5 ± 0.6, 8 hours post-uptake (Fig S1 M-O) indicating that 8 hours after phagocytosis, most phagosomes had not matured to phagolysosomes, since phago-lysosomal pH is expected to be pH < 5.5.

Together, these data indicate that non-immunostimulatory and ‘signal-free’ phagocytosed cargo mature stochastically, with fewer than 50% of internalized particles reaching lysosomes. These effects are independent of the method of analysis, lysosomal volumes, or the phagocytic capacity of the cell. To test whether these results were a feature of our designer phagocytic cargo, we repeated our studies with *E. coli* as our phagocytic cargo. Here, we observed that all phagocytosed bacteria were found in acidic phagolysosomes labelled by LTR at all time points tested post infection (Fig 1 F, G).

### LPS increases lysosomal localization of phagocytosed particles

To test our hypothesis that components from bacteria such as LPS promote lysosomal localization following phagocytosis, we activated RAW 264.7 cells by overnight treatment with LPS (18 hours at 100 ng/mL) and thereafter allowed particle uptake. We found that a significantly larger proportion of particles colocalized with lysosomes upon LPS-stimulation across all time points measured compared to untreated cells (control) (Fig 2 A, B). Similarly, in primary mouse peritoneal macrophages that were activated with LPS *ex vivo*, we observed that particles had a higher propensity to localize to the lysosomes than in cells that were not activated with LPS (Fig 2 C, D). While LPS activation is known to increase overall lysosomal volumes in macrophages, which we observe too, we found that the volume fraction of lysosomes in cells reduces upon activation because the volume of the cell itself increases (Fig S2 A). These data suggest that lysosomal volume changes are unlikely to influence the outcome of our image analysis.

**Figure 2:**
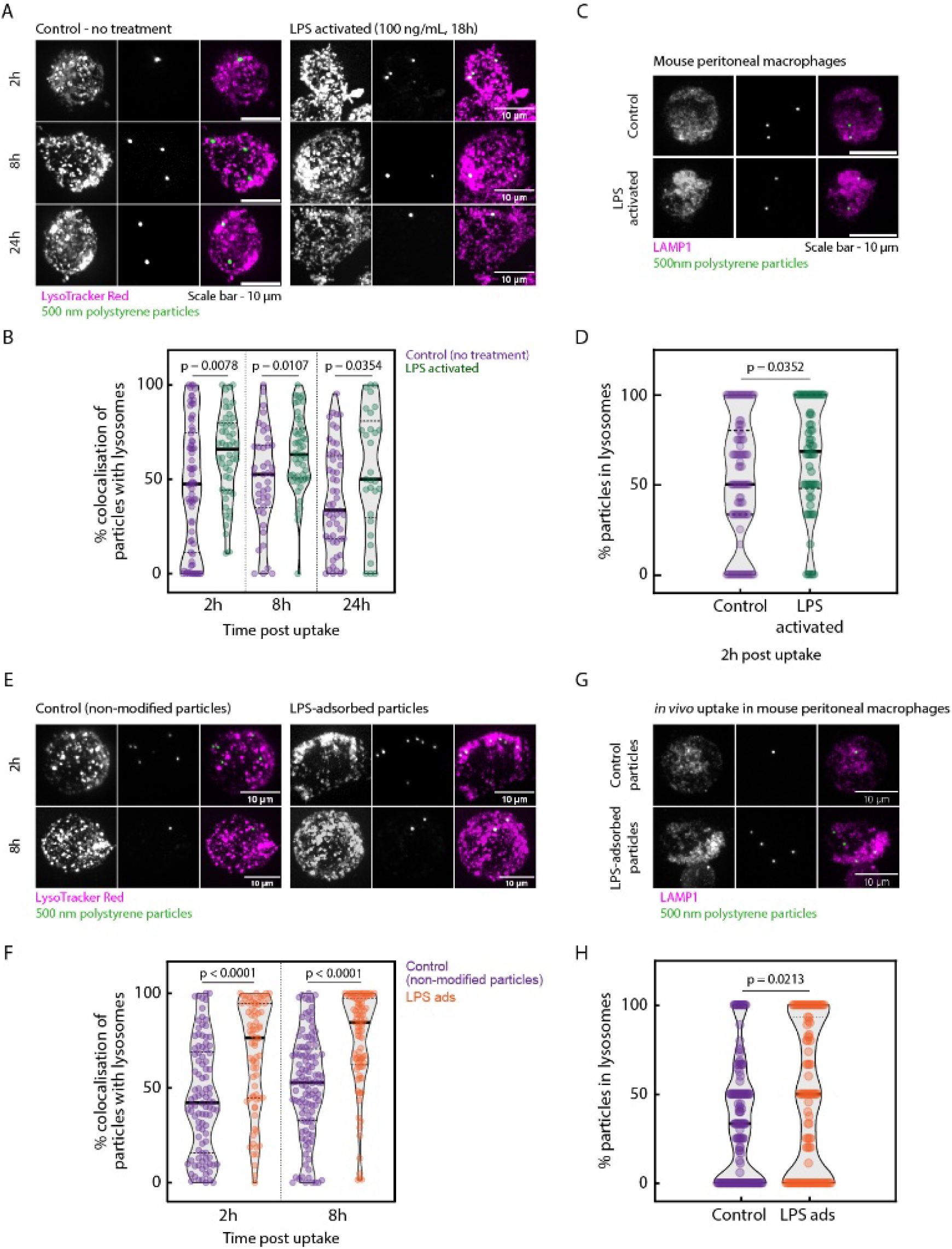
LPS enhances lysosomal localization of phagocytosed particles in RAW 264.7 cells and primary mouse macrophages. **A** – Representative MIPs of untreated cells (control) or LPS-activated RAW 264.7 cells at indicated time points post uptake of 500 nm non-modified polystyrene particles (Green) and stained with LTR (Magenta). Scale bar – 10 μm. **B –** Percentage colocalization of particles with LTR signal at indicated time points. Mann Whitney test; N = 3 independent experiments, 49 cells per group on average. **C –** Representative MIPs of primary peritoneal macrophages isolated from mice and either left untreated (control) or activated with LPS before *ex vivo* phagocytosis of particles. Magenta – LAMP1; Green – 500 nm-sized polystyrene particles. Scale bar – 10 μm. **D –** Percentage of internalized particles per cell that colocalize with LAMP1 in mouse peritoneal macrophages. Mann Whitney test; N = 3 independent experiments, 66 cells per group on average. **E –** Representative MIPs of RAW 264.7 cells after uptake of either control or LPS-adsorbed 500 nm-sized fluorescent particles at the indicated time points. Magenta – LysoTracker Red, Green – 500nm sized particles. Scale bars – 10 μm. **F –** Percentage colocalization of particles with LTR signal at indicated time points. Mann-Whitney test; N = 3 independent experiments, 94 cells per group on average. **G –** Representative MIPs of primary peritoneal macrophages isolated from mice after *in vivo* phagocytosis of particles. Magenta – LAMP1; Green – 500 nm sized particles Scale bars – 10 μm. **H –** Percentage of internalized particles per cell that colocalize with LAMP1 in mouse peritoneal macrophages. Mann Whitney test; N = 4 mice (control) and 3 mice (LPSads), 144 and 103 cells per group respectively. In all violin plots, dark lines represent median, and dashed lines represent quartiles of the data.

We then used a complementary approach, where we adsorbed LPS on to the particle surface and used LPS-adsorbed (LPS-ads) particles as phagocytic cargo. LPS adsorption was confirmed using fluorescent Alexa Fluor 488 – tagged LPS and imaging the particles (Fig S2 B, C). At both 2 h and 8 h post uptake, LPS adsorbed particles showed a higher propensity to localize to the lysosomes compared to control, uncoated particles (Fig 2 E, F). To test whether the effects of LPS signaling was physiologically significant, we injected either control (non-stimulatory, unlabeled particles) or LPS-ads particles into the peritoneal cavity of C57BL6 mice, and the peritoneal macrophages were isolated 2-hours post-injection. Further 3 hours post isolation, imaging the adhered primary cells revealed that LPS-ads particles had a higher tendency to localize to the lysosomes compared to uncoated control particles even *in vivo* (Fig 2 G, H).

Enhanced lysosomal localization of particles is also observed when non-modified, non-immunostimulatory particles are internalized along with either *E. coli* or LPS-ads particles in RAW 264.7 cells. That is, when the uncoated particles were introduced to cells along with either *E. coli* (Fig. S2 D-E) or LPS-ads (Fig. S2 F-G) particles in the same dish, a higher proportion of uncoated particles reached the lysosome, suggesting that the presence of a TLR-4 ligand in culture was sufficient to enhance phagosome maturation of all cargo. Together, these data demonstrate that LPS-induced signaling enhances phagosome – lysosome fusion, irrespective of the origin of the LPS signal.

### Additional ligands influence lysosomal localization of phagocytosed particles

Next, we aimed to delineate the effects of other ligands, given that our designer phagocytic assay is well suited to such reductionist approaches. We covalently conjugated either mouse immunoglobulin (IgG), bovine serum albumin (BSA) or folic acid to the surface of carboxylated polystyrene particles. We first confirmed that non-labelled carboxyl-functionalized particles had similar rates of delivery to the lysosome as non-modified particles (Fig S3 A, B). Then, we demonstrated that IgG and albumin conjugated particles showed enhanced localization to lysosomes whereas folic acid-conjugated particles showed no difference compared to unlabeled carboxyl-functionalized particles (Fig 3 A, B). Similar effects of IgG and folic acid-conjugated particles were observed in primary murine peritoneal macrophages in an *ex vivo* phagocytosis assay (Fig 3 C, D). Further, we tested the role of flagellin in promoting phagosome-lysosome fusion. We also tested the role of flagellin to test if other TLR-ligands promote phagosome – lysosome fusion. We find that pre-treatment with 250 ng/mL flagellin also enhances the lysosomal localization of particles 2h post uptake (Fig S3 C, D). These data prompted us to examine a potential common link between LPS, IgG, and albumin.

**Figure 3:**
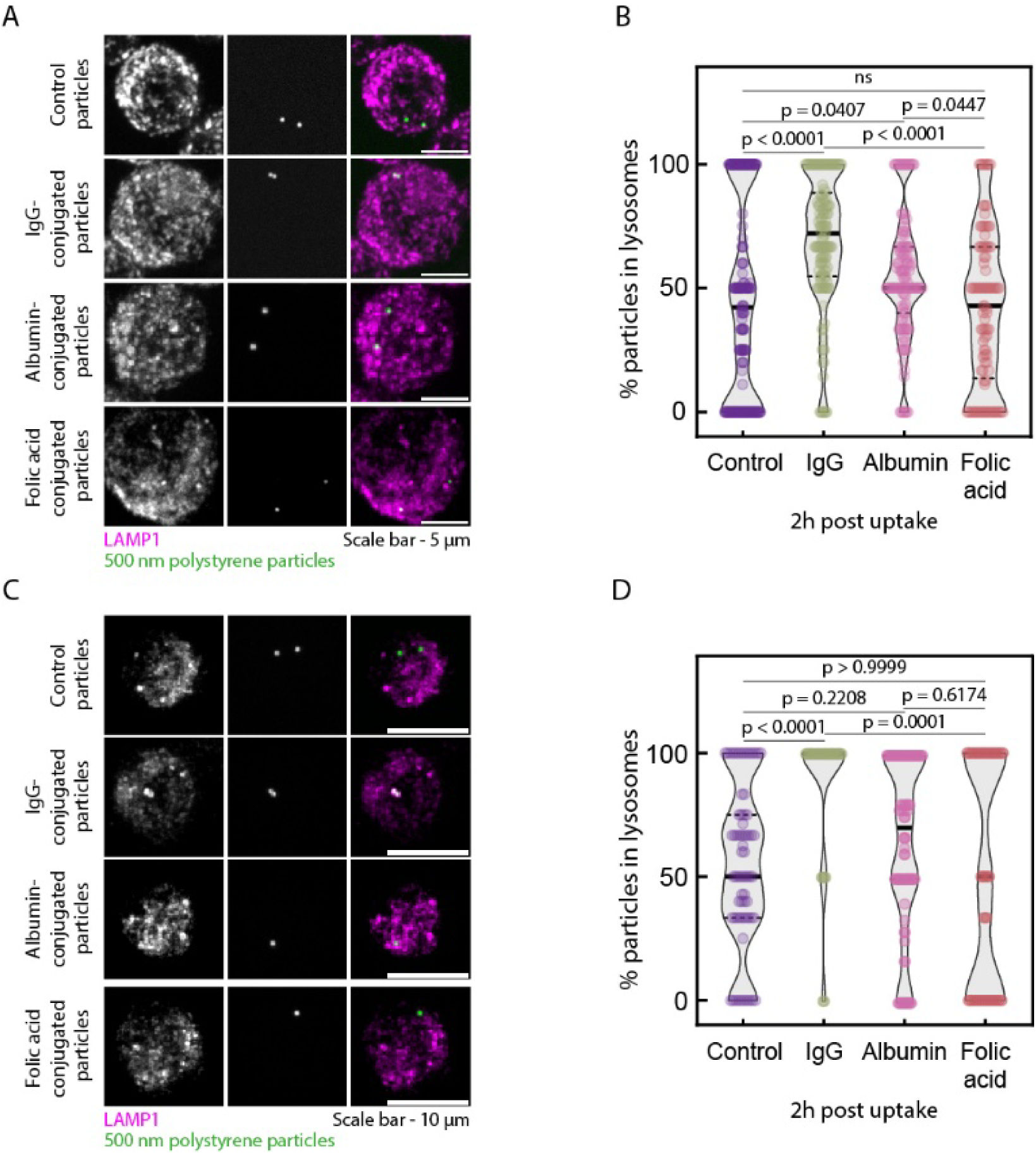
Conjugation of ligands to activate specific cognate receptors upon phagocytosis in RAW 264.7 cells and primary mouse macrophages. A – Representative MIPs of RAW 264.7 cells 2 hours after uptake of either control particles or particles conjugated with IgG, albumin or folic acid. Magenta – LAMP1, Green – particles. Scale bars – 5 μm. **B –** Percentage of phagocytosed particles per cell that colocalize with LAMP1 signal in RAW264.7 cells. Kruskal-Wallis test; N = 3 independent experiments, 130 cells per group on average. **C –** Representative MIPs of primary peritoneal macrophages isolated from mice 2h after *ex vivo* internalization of either control particles or particles conjugated with either mouse IgG, albumin or folic acid. Magenta – LAMP1; Green – 500 nm-sized polystyrene particles. Scale bars – 10 μm. **D –** Percentage of phagocytosed particles per cell that colocalize with LAMP1 signal in mouse peritoneal macrophages. Kruskal-Wallis test; N = 3 independent experiments, 58 cells per group on average. In all violin plots, dark lines represent median and dashed lines represent quartiles of the data.

### A potential role for p38 MAPK in LPS-induced phagosome maturation

LPS, IgG and albumin trigger TLR4 signaling, the Fc[R pathway and scavenger receptor A1 signaling, respectively. These three pathways converge at a stress-activated kinase – the p38α mitogen-activated protein kinase (henceforth, p38 MAPK) (Han et al., 1994; Nikolic et al., 2011; Ben et al., 2015; Coller and Paulnock, 2001; Linares-Alcántara and Mendlovic, 2022; Rose et al., 1997; Mkaddem et al., 2019). This enzyme has not been implicated in the folate-receptor signaling pathway (Chen et al., 2013; Nawaz and Kipreos, 2022). Further, flagellin is recognized by TLR5 on the cell surface and, like LPS, is also MyD88-dependent, and known to activate p38 MAPK (Hajam et al., 2017; Han et al., 1994). Hence, we tested whether p38 MAPK facilitates the maturation of phagosomes to phagolysosomes in the above scenarios. RAW 264.7 cells were treated either with the specific p38α MAPK inhibitor SB203580 or DMSO, the vehicle, for an hour before activation with LPS. Reduced p38 MAPK activation was confirmed by measuring IL-1β and TNF-α transcript levels which were respectively lowered and increased upon inhibitor treatment (Kim et al., 2004; Baldassare et al., 1999) (Fig. S4 A). We observed that upon inhibiting p38 MAPK the maturation of phagosomes to the lysosome dropped to levels observed with non-immunostimulatory particles in resting macrophages (Fig 4 A, B). Pre-treatment of RAW 264.7 cells with 10 µM SB203580 for 1 hour before uptake also reduced the enhanced lysosomal localization of LPS-ads particles (Fig 4 C, D).

**Figure 4:**
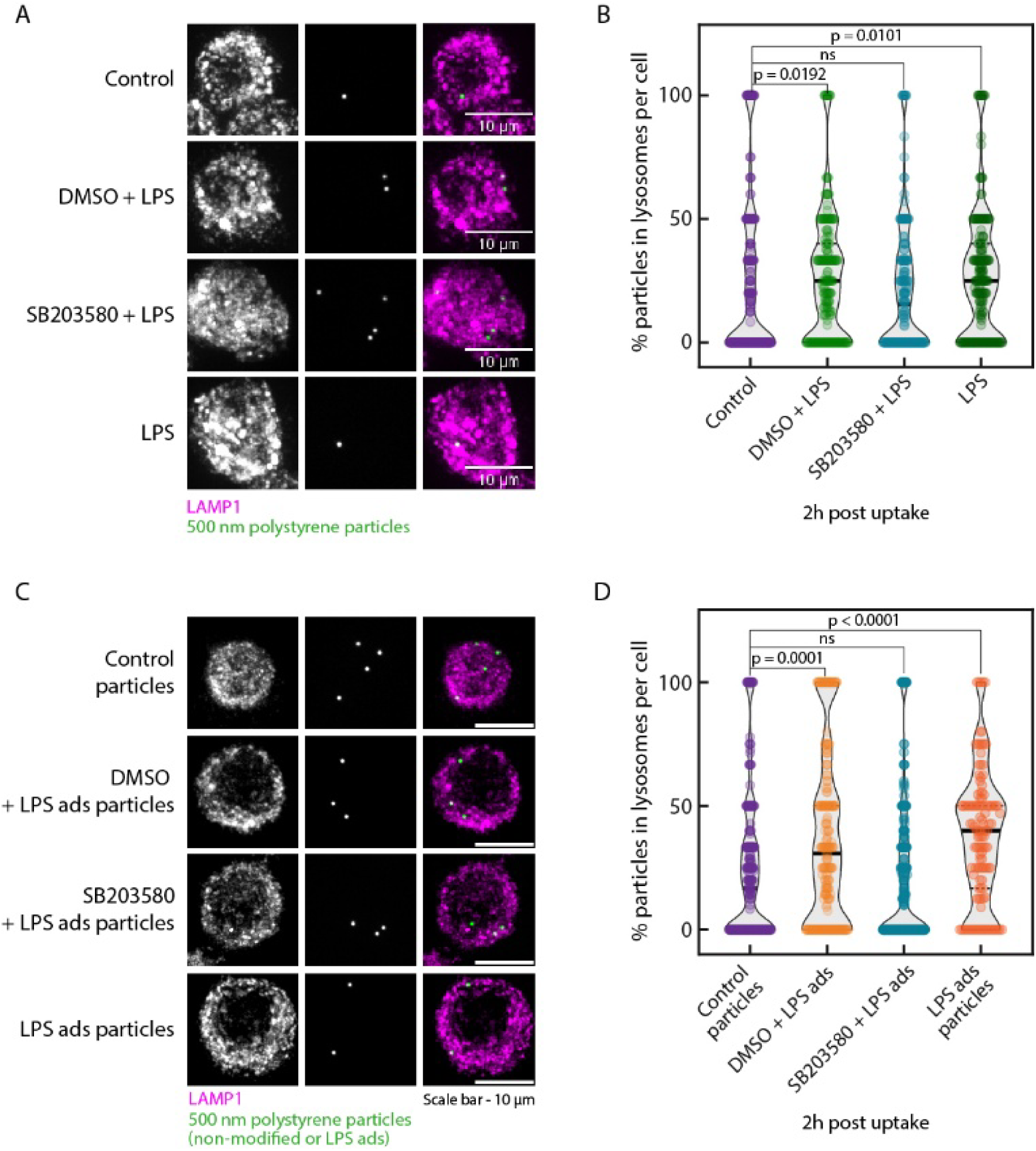
Inhibition of p38MAPK activity abrogates LPS - induced enhancement of phagosome maturation. **A –** Representative MIPs of RAW 264.7 cells 2 hours post uptake of particles. Cells were treated with p38MAPK inhibitor SB203580 (10 μM) or vehicle (DMSO) for 1h followed by 100 ng/mL LPS for 18h before uptake, or left untreated (control). Magenta – LAMP1, Green – 500 nm polystyrene particles. Scale bars – 10 μm. **B –** Percentage of particles taken up per cell that colocalize with LAMP1 signal 2 hours post uptake in RAW 264.7 cells after treatments described in A. Kruskal-Wallis (with Dunn’s post-hoc) test; N = 3 independent experiments, n = 219 cells per group on average. **C –** Representative MIPs of RAW 264.7 cells which were treated with either 10 μM SB203580 or vehicle (DMSO) or received no treatment before internalization of either non-modified (control) or LPS ads particles (all other groups). Treatments occurred 2 hours-post addition of particles. Magenta – LAMP1, Green – 500 nm polystyrene particles (either non-modified – control or LPS adsorbed). Scale bars – 10 μm. **D –** Percentage of particles taken up per cell that localize to lysosomes, 2h post uptake after treatments described in C. Kruskal-Wallis (with Dunn’s post-hoc) test; N ≥ 3 independent experiments, n = 226 cells on average per group. Data are plotted as violin plots with black lines indicating median, dashed lines indicating quartiles, and colored lines indicating mean of the data.

To further probe the role of p38 MAPK in enhancing phagosome maturation, we reduced p38 MAPK levels in RAW 264.7 cells using RNA interference and achieved a modest reduction of ∼ 40% (Fig S4 B). With this level of knockdown, our automated analysis methods did not reveal a significant difference in lysosomal localization of particles compared to un-transfected and scrambled-siRNA controls (Fig S4 C). When activated with LPS, there is an insignificant trend for reduced lysosomal localization of particles in siRNA-transfected cells (Fig S4 D). However, a knockout-model or more efficient knockdown of p38 MAPK would be more informative to definitively test the role of p38 MAPK in regulating phago-lysosomal fusion. Nevertheless, the data support the finding that phagosome maturation is stochastic in the absence of stimulatory signals.

### Phagosomal pH dynamics differ with LPS treatment and depend on p38 MAPK activity

Next, we explored whether along with enhanced lysosomal localization under LPS-activation, the phagosome acidification rates are affected. We used a small molecule pH sensor, derived from a pentacyclic pyrilium fluorophore PS-OH denoted SeRapHin (self-ratiometric pH indicator; pKa = 7.4 and 8.6), which possesses two emission spectra (SeRapHin green – Exc: 445 nm, Em: 500-550 nm; SeRapHin red – Exc: 488 nm, Em: 625-700 nm) and its ratio (green/red) is inversely proportional to pH(Chakraborty et al., 2020).

We conjugated SeRapHin to the particle surface to make pH-reporting SeRapHin-beads. A standard curve was created by clamping RAW 264.7 cells that had internalised these beads, in buffers of varying pH supplemented with ionophores nigericin and monensin or valinomycin (Fig S5 A – C). The photostability of SeRapHin-beads and fidelity of reporting pH was validated by imaging SeRapHin-beads in extracellular medium over time as a control while performing our time-lapse experiments, described further below (Fig S5 D).

By monitoring pH changes in the early stages of phagosome maturation, our data indicate that early decisions post uptake determine the maturation fate of a phagocytosed cargo. Through time-lapse microscopy, we find that LPS-treated cells have consistently more acidic phagosomes compared to control, and this difference is lost if the cells are pre-treated with SB203580 (Fig 5 A, B). In fact, we find that phagosomal fate (measured as phagosomal pH) is determined early, within first 10 – 15 minutes of uptake, in the phagosome maturation process. Together, these data suggest that specific cell surface protein-ligand interactions which activate p38 MAPK, leads to an increased and faster lysosomal localization of phagocytosed cargo, and that inhibiting p38 MAPK activity reduces the lysosomal delivery of phagocytosed cargo.

**Figure 5:**
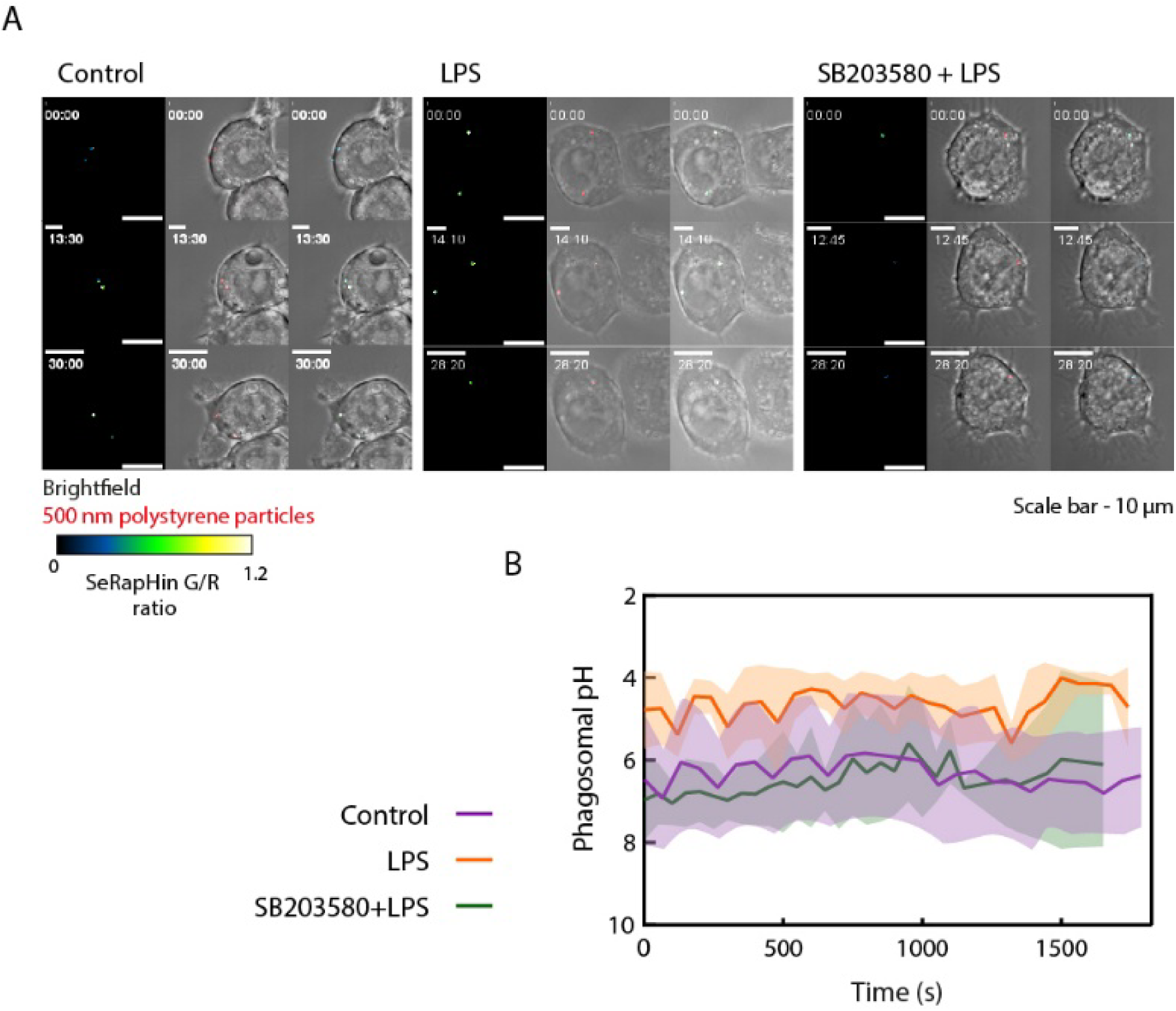
Phagosomal acidification is also enhanced upon LPS-activation and depends on p38 MAPK activity. **A** – Representative MIPs of Z-stack time lapses acquired of SeRapHin-conjugated polystyrene beads internalized by RAW 264.7 cells that were left untreated, or activated with LPS, or treated with SB203580 prior to LPS activation. Grayscale – brightfield, Red – 500 nm polystyrene particles, Green Fire Blue LUT – SeRapHin G/R ratio masked for visualization of signal from particles alone. Scale bars indicated on the bottom right are 10 μm and time stamps are indicated on the top left in mm:ss. **B –** Average pH as reported by SeRapHin-conjugated 500 nm polystyrene particles. Purple – control cells, Orange – Cells treated with LPS for 18h, Green – Cells treated with SB203580 for 1 hour prior to LPS treatment. n = 15, 10 and 13 trajectories respectively across N > 3 independent experiments. Colored lines and shaded regions respectively indicate the mean pH and standard deviation across the trajectories.

## Discussion

In macrophages, phagosome maturation resulting in phago-lysosomal fusion is considered *de facto* unless the phagocytosed cargo has specific mechanisms of stalling or escaping the process (Simeone et al., 2021; Vergne et al., 2003; Ahmad et al., 2024). Previous studies provide evidence of polystyrene particles not reaching lysosomes in both professional and non-professional phagocytes, but it was not explored further (Oh and Swanson, 1996; Rejman et al., 2004; De Chastellier et al., 1995; de Chastellier and Thilo, 1997). We therefore leveraged these findings to create a precise and tunable phagocytic assay based on polystyrene particles as model phagocytic cargo to test the role of liganding specific cell-surface receptors in phagosome maturation.

One significant observation is the variability of phagosome maturation in macrophages. For example, we observe that in some cells phagocytosed particles localizing to the lysosome in an all-or-none fashion, suggesting heterogeneity in cellular behavior with respect to phagosome maturation. However, in other cells cultured in the same dish, we observed that ∼50% of internalized particles mature to lysosomes, indicating that differential maturation rates may be observed in the same cell, as has been reported earlier for specific phagosomal markers (Henry et al., 2004). Signaling triggered by ligands such as LPS, albumin, IgG and possibly other TLR-ligands (flagellin), shifts the distribution towards most particles maturing to lysosomes.

Particles coated with LPS (LPS-ads) display enhanced lysosomal localization even within 2 hours after uptake in cells, indicating that early events triggered after LPS-TLR4 interaction are sufficient to determine the fate of these phagosomes. These results contradict another study of phagosome maturation in macrophages, where no difference in acidification rates were seen when the phagocytic cargo used were silica beads coated with either IgG, mannosylated-albumin with or without an LPS coating (Yates and Russell, 2005). These differences may arise from the fact that silica is known to be immunostimulatory and cause cellular changes in macrophages (Kim et al., 2012). Hence, it is possible that adding LPS on top of these signals does not further enhance maturation. Our phagocytic cargo are polystyrene beads that are not known to stimulate macrophages and is therefore ideal to test the downstream contribution of stimulating cell surface receptors to phagosome maturation.

We identified p38 MAPK as a signaling node for the various cell surface receptors that enhance phagosome maturation. Chemical inhibition of p38 MAPK reduced LPS-induced enhancement of phagosome maturation. Along with increased delivery to the lysosome, we find phagosomal acidification kinetics also differ in LPS-activated macrophages by using a self-ratiometric pH reporter conjugated to the particles. A few studies have reported phagosomal pH is ≤6 within 10-15 minutes of cargo uptake (Yates et al., 2005; Canton et al., 2014; Thekkan et al., 2019), and through our live-imaging experiments we observe a discernible difference in pH across LPS-treated and control cells within 10 minutes of particle uptake. Phagosomes in LPS-treated cells are consistently more acidic than the latter on average, and this difference is lost when p38 MAPK activity is inhibited. Phagosomal pH depends on several factors, such as V-ATPase abundance and activity, phagolysosomal fusion events and numbers and function of ion channels on the phagosomal membrane. LPS enhances phago-lysosome fusion in bone-marrow derived macrophages as early as 5 minutes post cargo uptake, and can explain the observed enhanced acidification kinetics triggered by LPS (Yates and Russell, 2005).

While siRNA-mediated knockdown of p38 MAPK resulted in a modest but significant reduction of the protein levels (∼40%), it failed to show a significant difference in rates of phagosome maturation of particles upon LPS-activation. Thus, either knockdown levels of p38 MAPK are insufficient or, in an unlikely scenario, knockdown causes compensatory changes in the cell, which are not seen upon acutely perturbing p38 MAPK with a chemical inhibitor. A knockout model could be employed in future studies to establish the role of p38 MAPK in maturation.

p38 MAPK has previously been implicated in Rab5 membrane-cytosol cycling to promote endosome maturation (Cavalli et al., 2001). Also, p38 MAPK inhibition has previously been shown to inhibit phagosome maturation, but specifically in the context of TLR-4 signalling (Blander and Medzhitov, 2004). Our results build on these previous results and suggest that ligands interacting with cell surface receptors that signal via p38 MAPK, which include LPS, flagellin, IgG and albumin, increase both the amount and rate of phagosome maturation. Together, these findings show that in the absence of any external signals, phagosome maturation is stochastic and is enhanced by the liganding of specific cell surface receptors that converge on the p38 MAPK signalling pathway. Understanding how phagosome maturation is regulated will help us better evaluate its inherent heterogeneity and thereby aid in designing targeted drugs, vaccines or diagnostic particulates.

In summary, we have demonstrated that maturation of phagosomes is independent of the phagocytic capacity of the macrophage. Rather it is a stochastic phenomenon, wherein fewer than 50% of internalised particles per cell undergo fusion with lysosomes. Specific signals on the other hand can enhance phagosome-lysosome fusion, and these signals include LPS or flagellin in solution and LPS, IgG or albumin conjugated on the surface of the particle, all of which ligand various cell surface receptors. Additionally, LPS-induced signaling enhances acidification rates of the phagosome. These effects are lost upon pharmacologically inhibiting p38 MAPK, suggesting a potential role for the enzyme in regulating phagosome maturation in macrophages.

## Materials and Methods

### Materials

Spherical polystyrene particles (either non-modified or carboxyl-terminated surfaces) of 500 nm diameter and labelled fluorescently (excitation/emission maxima of either 480/520 nm or 660/690 nm, Bangs Laboratories, Indiana, USA) were used as phagocytic targets. Lysosomes were stained using LysoTracker Red DND-99 (Invitrogen, L7528). For immunofluorescence assays, F-actin was stained with Rhodamine Phalloidin (Invitrogen, USA or abcam, UK) or Actin Geen (Invitrogen, USA), nucleus was stained with Hoechst 33342 (Invitrogen, USA). Primary antibody against LAMP1 (rat anti-mouse) was procured from Developmental Studies Hybridoma Bank, Iowa (Clone number: 1D4B) and primary and secondary antibodies used for Western blot assays (rabbit anti-mouse: p38 MAPK – polyclonal, GAPDH – clone 14C10 and β tubulin – polyclonal, and goat anti-rabbit: HRP-linked anti-rabbit IgG) were purchased from Cell Signaling Technologies, USA.

### Cell culture and treatments

RAW 264.7 cells (Sigma Aldrich, now Merck, USA) were cultured in Dulbecco’s Modified Eagle’s Medium (DMEM) (Cell clone, Genetix Biotech, India and MP Biomedicals, USA) supplemented with 10% fetal bovine serum and 1% anti-bacterial and anti-mycotic agent (Thermo Fisher Scientific, USA). For phagocytosis studies, cells were seeded at a density of ∼0.5 x 10^5^ / cm^2^ on gelatin-coated, acid-cleaned 12 mm no. 1 glass cover slips (Blue Star, India) or gelatin-coated glass-bottomed imaging dishes (ibidi, Germany). Particles were incubated with macrophages for 1 hour at a cell to particles ratio of 1:200, which we defined as the pulse period. At the end of the pulse period, particles that were not internalized were removed with multiple, thorough washes with 1X PBS. Thereafter, cells were supplemented with fresh complete medium and taken for further processing at different time points as specified.

Cells were treated with the following ligands 18 hours prior to uptake of particles, by adding them to the culture media in the following concentrations – 100 ng/mL lipopolysaccharide (from *E. coli* O111:B4, Sigma) and 250 ng/mL flagellin (Sigma Aldrich). The compounds were maintained in solution throughout the pulse and chase periods as well. Cells were treated with 10 µM SB203580 for 1 hour prior to LPS treatments (or introduction of LPS-ads particles) to inhibit p38 MAPK activity. Equivalent volume of DMSO (final concentration of DMSO in media not exceeding 0.1%) was added before LPS treatment as a vehicle control. The drug or DMSO and LPS were maintained in the media during the pulse and chase periods as well.

### *E. coli* infection study

Constitutively fluorescent bacteria were prepared by transforming DH10β cells (a kind gift from Prof. Sandhya Visweswaraiah, IISc) by electroporation with the pDiGc plasmid (Addgene), which ensured constitutive EGFP-expression. These were incubated with RAW 264.7 macrophages for 1 hour at a multiplicity of infection 10:1 in antibiotic-free, serum-supplemented DMEM, after which bacteria which were not internalized were washed off with 1X PBS and fresh, complete medium supplemented with 100 µg/mL gentamycin (Sigma Aldrich) was introduced to the cells. After 1 hour, gentamycin concentration in the wells was reduced to 25 µg/mL.

### Labelling of lysosomes

Macrophage lysosomes were stained using with LysoTracker Red DND-99 (LTR) (Thermo Fisher Scientific, USA), or by immunostaining for lysosomal-associated membrane protein (LAMP1). For the former, macrophages were stained with 100 nM LTR for 10 minutes at 37°C, 5% CO2 in live-cell imaging buffer (20 mM HEPES, 140 mM NaCl, 2.5 mM KCl, 1.8 mM CaCl2, 1 mM MgCl2, 4 mg/mL glucose, pH 7.4). Thereafter, the staining solution was washed off and cells were immediately imaged live in the live-cell imaging buffer.

For immunofluorescence of LAMP1, cells growing on gelatin-coated coverslips were fixed at required time points using freshly prepared 4% paraformaldehyde and permeabilized using 0.1% Triton-X 100. Blocking was carried out prior to each antibody-incubation step with 5% bovine serum albumin (BSA) solution for 60 minutes at room temperature. Primary antibody incubation was carried out with antibody diluted at 1:100 in 2% BSA solution and incubating the cells overnight (16h – 18h) at 4°C. Secondary antibody staining was carried out at room temperature for 1 hour with a secondary-staining mix which constituted the secondary antibody at 4 µg/mL, rhodamine phalloidin (Abcam and Thermo-Fisher Scientific, USA) at concentrations recommended by each manufacturer and Hoechst 33342 (Thermo-Fisher Scientific, USA) at 2 µg/mL, in 2% BSA solution. All incubations were preceded and followed with thorough washes in 1X PBS and all mentioned solutions used 1X PBS as the solvent. Coverslips were finally mounted on glass slides (Blue Star, India) with ProLong Diamond antifade mountant (Thermo Fisher Scientific, USA), which was allowed to cure overnight at room temperature.

### Covalent conjugation of ligands

Mouse immunoglobulin, bovine serum albumin, folic acid, ethylenediamine (Sigma Aldrich, now Merck, USA), and the self-ratiometric pH indicator SeRapHin were conjugated on to fluorescent polystyrene particles that had free carboxyl groups on their surface by carbodiimide crosslinking as described below. Carboxyl groups were activated with 40mM 1-Ethyl-3-[3-dimethylaminopropyl]carbodiimide hydrochloride (EDC) and 50 mM N-hydrosuccininmide (NHS) prepared in 100mM 2-(N-morpholino)ethanesulfonic acid buffer (pH 6), following which the particles were washed with 1X PBS and incubated on a rotor-shaker with the ligand solution (prepared in PBS) for 3 hours to allow for conjugation. Unbound ligands were removed by four washes with 1X PBS at the end of the incubation period. For folic acid conjugation, since the pterin moiety consisting of the pyrazino[2,3-d]pyrimidine nucleus is recognized by the folate receptor, the particles were first conjugated with ethylenediamine and then the exposed amine groups on the particle surface were conjugated with the glutamate moiety of folic acid following a similar protocol as mentioned above.

The amine-group surface-modified particles were also used to conjugate pHrodo Red succinimidyl ester (henceforth pHrodo). Amine-modified particles were resuspended in 100 mM carbonate-bicarbonate buffer (0.1M Na_2_CO_3_, 0.1M NaHCO_3_, pH 8.4) with 0.15 mM pHrodo and the reaction was allowed to proceed for 1 hour at room temperature. Unbound pHrodo was washed off by multiple spins with 1X PBS. Conjugation was confirmed by flow cytometry.

For all experiments with conjugated particles, corresponding particles used for the ‘control’ group denote particles which were subjected to the same reaction conditions and washes, and received 1X PBS instead of the ligand to be conjugated.

### Adsorption of LPS

Non-modified or carboxylated fluorescent, polystyrene particles were washed in 1X PBS and resuspended in 500 µg/mL solution of lipopolysaccharide in 1X PBS. This mix was agitated by placing the solution on a rotor shaker for 3h, after which free LPS in the solution was removed through multiple washes with 1X PBS in a microcentrifuge. Surface modification of particles following adsorption was confirmed by using fluorescent AlexaFluor 488-conjugated LPS (Thermo-Fisher Scientific, USA) and observing the LPS-adsorbed particles under a fluorescence microscope.

### Animal studies

All animal studies were conducted in accordance with the Control and Supervision Rules, 1998 of the Ministry of Environment and Forest Act (Government of India), and the Institutional Animal Ethics Committee, IISc. Experiments were approved by the Committee for Purpose and Control and Supervision of Experiments on Animals (permit number CAF/ethics/718/2019). Female, healthy and specific-pathogen free C57BL6 mice (acquired from Hylasco Biotechnology Hyderabad India a subsidiary of Charles River Laboratories) that were 8-12 weeks old, were injected with a 200 µL solution of sterile saline containing 6 x 10^6^ 500 nm-sized fluorescent particles. After 2 hours, the mice were euthanized and peritoneal exudate collected by flushing with sterile 1X PBS-EDTA. Cells were spun down and resuspended in complete DMEM and allowed to adhere to culture plates or gelatin-coated cover slips for 3 hours after which the floating cells were washed off and the adhered cells were either fixed for immunofluorescence or scraped for characterization by flow cytometry.

For *ex vivo* uptake studies, 8-12 weeks old, female, healthy C57BL6 mice were euthanized, and peritoneal exudate was collected in a similar fashion as described above. Peritoneal cells were seeded in culture plates or gelatin-coated cover slips and allowed to adhere for 3 hours, after which floating and loosely adhered cells were washed off. The remaining adhered cells were ∼ 80 % macrophages (as characterized by flow cytometry of cells scraped at the time point) and were allowed to internalize particles following the same protocol as described earlier for RAW264.7 macrophages.

### pH measurements using pHrodo and SeRapHin

pHrodo Red-conjugated particles were incubated with RAW264.7 cells for 1 hour, and the cells were then clamped for 60 minutes in the following buffers supplemented with ionophores nigericin and valinomycin at 10 µM each – 140 mM KCl, 1 mM MgCl2, 1 mM CaCl2, 5 mM glucose buffered with 0.1 M acetic acid and 0.1 M sodium acetate (for pH 4 and pH 5) or 0.1M MES (for pH 6 and pH 7) to create a standard curve between pH and pHrodo intensities.

For clamping of SeRapHin-conjugated particle-containing phagsosomes, RAW 264.7 cells that had internalized SeRapHin-conjugated particles were first mildly fixed in freshly prepared 2.5% paraformaldehyde for 5 minutes and then washed with 1X PBS. Thereafter, cells were clamped for 30-40 minutes with ionophore (either 10 µM nigericin and 10 µM monensin, or 10 µM nigericin and 10 µM valinomycin) supplemented UB4 (20mM HEPES, 20mM MES, and 20mM sodium acetate, 140 mM KCl, 5 mM NaCl, 1mM MgCl2 and 1mM CaCl2), with pH of the buffers rising in increments of 0.5 starting from pH 4 up to pH 7.5. SeRapHin Green intensity is defined as the emission collected in the 500 nm – 550 nm band when excited at 445 nm, SeRapHin Red intensity is defined as the emission collected in the 660 – 750 nm band when excited at 488 nm. SeRapHin Green/Red (G/R) ratios are a readout for pH and the ratio is inversely proportional to pH of the solution.

For acquiring pH trajectories of phagosomes, cells growing in glass-bottomed dishes were washed and immersed in live-cell imaging buffer (described earlier) and incubated with SeRapHin-conjugated particles for 5 minutes at 4°C before taking the dish for imaging on a heated-stage (37°C) to synchronize particle uptake. To optimize tracking, Z-stacks of multiple selected fields in a dish were acquired continuously with 0 or minimum possible delay. Details of microscopy and quantification of these data are further described under microscopy and image analysis.

### Microscopy

Unless specified otherwise, all immunofluorescence images (and images of LysoTracker-stained cells) were acquired on an inverted microscope (Ti2E, Nikon) with a spinning-disk module (CSU-X1, Yokogawa) and an EMCCD camera (iXon Ultra-897, Andor). Solid-state lasers at wavelengths 405 nm, 488 nm, 561 nm and 640 nm (iChrome Multi-Laser Engine, Toptica) were used as the excitation source. Corresponding band-pass emission filters (B460/80, B520/35, B617/73, and B685/40) as fitted on the CSU-X1 were utilized. A CFI Plan Apochromat VC 100XH (NA 1.40, WD 0.13 mm) oil-immersion objective (Nikon) was used to acquire Z-stacks while maintaining a step-size of 0.3 µm to capture the entire cell. Z-stacks in live cells were acquired with channels alternating with every slice to maximize accuracy of lysosomal and particle localization inside the cell. For live cells, temperature was maintained at 37°C during imaging by using a stage top incubator (Okolab, Italy).

Some immunofluorescence images (for data represented in Fig S5 C, D) were acquired on an inverted microscope (DMi8, Leica) fitted with a spinning-disk (Dragonfly 400, Andor) and equipped with an sCMOS detector (Sona, Andor) using a 100X oil-immersion, 1.4 NA objective (Leica). Excitation was performed through laser lines 405 nm, 488 nm, 561 nm and 637 nm (High Power Laser Engine, Andor) and emission collected on the corresponding filter sets as fitted on the Dragonfly spinning-disk unit. The PCC cutoff for this modality was accordingly modified to 0.6 (described under image analysis).

Time-lapses with SeRapHin-beads and corresponding clamping experiments were acquired either on the Stellaris 8 (Leica) or the LSM 980 (Zeiss) and the specifications for each setup are provided below.

Stellaris 8 (Leica), an inverted, laser-scanning confocal microscope fitted with a stage and objective heater to maintain the temperature of the sample as well as the objective was utilized. Images were acquired using an HC Plan Apochromat 63X CS2 (NA – 1.40, WD – 0.14 mm) oil-immersion objective (#506350, Leica). The tunable white light laser available on the system was used for simultaneous excitation of SeRapHin at 448 nm (generated through the acousto-optical beam splitter) and 488 nm (notch generated from traditional, beam-splitting mirrors) and the respective emissions were collected in the ranges 500 nm – 550 nm (titled ‘SeRapHin Green’) and 660 nm – 750 nm (titled ’SeRapHin Red’) on the hybrid photodiode detectors HyD S and HyD R respectively, each set to the continuous mode. Particles were inherently fluorescent in the far-red channel and were detected by exciting at 635 nm with the emission collected on 642 nm – 660 nm on the HyD X detector together with a transmitted brightfield image collected on a photomultiplier tube detector. SeRapHin-conjugated particles were tracked in cells by acquiring time lapses of multiple Z-stacks that were acquired with a step size of 0.4 µm and minimized time intervals (averaging out to 60-80 second time intervals across experiments). For calibration, Z-stacks of SeRapHin-particles either *in vitro* or in clamped cells were acquired with the same imaging parameters used for tracking.

The LSM 980 (Zeiss) is also an inverted, laser-scanning confocal microscope and a Plan-apochromat 63X oil-immersion objective (Zeiss) was used for imaging. Laser lines at 445 nm and 488 nm wavelengths were used to excite SeRapHin and emissions were collected at 500 – 550 nm and 685 – 735 nm respectively on the GaAsP spectral detectors (Zeiss) fitted with the microscope. All other imaging and tracking parameters were kept the same as mentioned above.

### Image analysis

All automated quantification of microscopy data was performed using custom scripts written in IJM on Fiji, and MATLAB, and the scripts have been provided on https://github.com/Immunoengineeringlab/phagosomeMaturation-imageAnalysis-MS. The rationale for each of the object-overlap-based methods and Pearson’s correlation coefficient-based methods to quantify colocalization of particles with lysosomes is described below.

For cells where lysosomal lumen was stained using LTR, an object overlap based method was used to quantify colocalization. Lysosomal and particle signals were thresholded from cropped Z-stacks of individual cells using the Otsu algorithm on a difference of Gaussians processed (radii: 1 and 2 pixels) lysosome signal stack and the Intermodes algorithm on the particles stack on Fiji. The fraction of particle voxels that overlapped with lysosome voxels was read as *percentage of colocalisation of particles with lysosomes* in the cell. For cells where lysosomal membrane was stained (via immunofluorescence for LAMP1), a modified Pearson’s correlation coefficient (PCC) based method was used to quantify colocalization. Briefly, in a cropped Z-stack that encompassed a single cell alone, mean intensities across the Z-stack were recorded for LAMP1 and particle channels in individual particle ROIs. The ROIs were determined automatically from maximum intensity projections of the particle channel, therefore cells where even one aggregate of particles appeared in the projected image were excluded from the analysis pipeline. The PCC between LAMP1 and particle signal was calculated for these recorded intensities for each particle inside the cell, and based on manual checking, a cut-off of 0.7 for the PCC was determined to classify whether the particle would be identified as localizing withing the lysosome or not. Thus, *the percentage of particles in lysosomes per cell* were quantified. Cutoff for the Pearson’s correlation coefficient to identify a particle as being co-localized with lysosome signal was taken to be 0.5 for *ex vivo* uptake experiments in primary macrophages. Images were also verified for colocalization extents by visual inspection.

For quantifying pH trajectories in SeRapHin experiments, we extracted SeRapHin G/R ratios as follows. Maximum intensity projections (MIP) of Z-stacks were used to identify particle ROIs in both the calibration and tracking assays, and from these MIPs, the mean SeRapHin Red and Green emission intensities were recorded after subtracting the mode of each projected channel from the respective images to subtract background intensity. For calibration, the ratio of mean green to red intensity from thus background-subtracted images were plotted and the mean G/R value thus obtained at each pH was used to fit to a linear regression. G/R values obtained from tracking experiments were then interpolated from each experiment’s calibration curve and a rolling average of 3 time points was plotted to obtain the pH trajectories.

For pHrodo experiments, pHrodo intensity was recorded from the slice where the particle was in focus (determined by slice where maximum fluorescence intensity recorded from the particle in the Z-stack) and calibration was performed with these mean intensity values across four pH values 4, 5, 6 and 7.

### RT-qPCR assay

RAW 264.7 cells were seeded to ∼75% confluence in 12-well plates and allowed to adhere overnight (18 hours) with or without 100 ng/mL LPS. After incubation, cells were lysed with 1 mL of TRI reagent (Sigma Aldrich) and intracellular RNA was isolated from these lysates using RNeasy Mini Kits (Qiagen, USA) which included an on-column DNAse digestion step. 1 µg of RNA from each sample was reverse-transcribed to complementary DNA using the iScript cDNA synthesis kit (BioRad, USA) as per the manufacturer’s recommendations. The synthesised cDNA was analysed by amplification through a real time quantitative polymerase chain reaction (RT-qPCR), using TB Green Premix Ex Taq I (Takara Bio Inc., Japan) normalised to the expression levels of the house-keeping gene glyceraldehyde 3-phosphate dehydrogenase (*Gapdh*). The forward and reverse primer sequences for IL 1β were 5’-CAACCAACAAGTGATATTCTCCATG-3’ 5’ GATCCACACTCTCCAGCTGCA-3’ respectively, while for TNF-α they were 5’ GTGCCTATGTCTCAGCCTCTT-3’ and 5’-GCCATAGAACTGATGAGAGGGAG-3’ respectively.

### RNA interference – transfection of siRNA

The sense strands of the oligonucleotides used for RNA interference - based knockdown of p38 MAPK are as follows: 5’ GAU-GAA-CUU-CGC-AAA-UGU-A)TT 3’ and 5’ (CAC-UGA-AUC-CAG-UGU-CAA-U)TT 3’ and scramble siRNA (Eurogentec, Belgium).

RAW 264.7 cells were transfected with these oligos at 25 nM 24 hours after seeding using the INTERFERin transfection reagent (Polyplus, France) and optiMEM (Thermo Fisher Scientific, USA) according to the manufacturer’s recommendations. Cells were washed and media replenished 6 hours after transfection, and were taken for further processing another 42 hours later. Silencing of p38 MAPK was tested with a western blot assay.

### Western blot assay

For western blot assays, cells were lysed in mammalian cell lysis buffer (Gold Biotechnology, Inc., USA) supplemented with phosphatase and protease inhibitor cocktail (Sigma), and soluble proteins were collected by centrifuging these lysates. After determining protein concentrations using Bradford’s assay, cell lysates containing 10 µg total protein along with a standard protein ladder (Precision Plus Protein Dual Colour Standards, BioRad) were run on a 10% resolving gel through sodium dodecyl sulphate-polyacrylamide gel electrophoresis and transferred onto a polyvinylidene fluoride membrane by wet transfer at constant current. A ponceau stain was done to confirm transfer of proteins and the blot was then washed and blocked with 5% BSA solution prepared in tris-buffered saline and tween buffer. Primary staining was performed overnight at 4 °C on a rocker and secondary staining was performed at room temperature for 60 – 90 minutes on a rocker. A 1:1 mix of peroxide and luminol reagents (Clarity Western ECL Substrate, BioRad) were added to the blot just before imaging to develop signal from HRP-conjugated secondary antibody. Blots were imaged on a ChemiDoc MP imaging system (BioRad) and images quantified on ImageLab software (BioRad).

### Statistics

Comparisons across groups were done using the Mann-Whitney Test or the Kruskal-Wallis test with Dunn’s Multiple Comparisons post-hoc test when required. GraphPad Prism 8.0 was used to perform each of these tests and graphs were plotted on Prism GraphPad 8.0 and MATLAB R2022a (MathWorks) and figures were edited and assembled on Adobe Illustrator 2019.

## Acknowledgments

We thank the Biosciences Imaging Facility at the Indian Institute of Science, the Central Imaging & Flow Cytometry Facility at the National Centre for Biological Sciences, India and the Integrated Light Microscopy Core at The University of Chicago, USA for their facilities. We also acknowledge the support provided by the Central Animal Facility, Indian Institute of Science.

## Data and materials availability

All data are available in the main text or the supplementary materials.

## References

Ahmad, A., J.M. Khan, B.A. Paray, K. Rashid, and A. Parvez. 2024. Endolysosomal trapping of therapeutics and endosomal escape strategies. Drug Discovery Today. 29:104070. 10.1016/j.drudis.2024.104070.

Baldassare, J.J., Y. Bi, and C.J. Bellone. 1999. The Role of p38 Mitogen-Activated Protein Kinase in IL-1β Transcription1. The Journal of Immunology. 162:5367–5373. doi:10.4049/jimmunol.162.9.5367.

Ben, J., X. Zhu, H. Zhang, and Q. Chen. 2015. Class A1 scavenger receptors in cardiovascular diseases. British Journal of Pharmacology. 172:5523–5530. doi:10.1111/bph.13105.

Blander, J.M., and R. Medzhitov. 2004. Regulation of Phagosome Maturation by Signals from Toll-Like Receptors. Science. doi:10.1126/science.1096158.

Canton, J., R. Khezri, M. Glogauer, and S. Grinstein. 2014. Contrasting phagosome pH regulation and maturation in human M1 and M2 macrophages. Molecular Biology of the Cell. doi:10.1091/mbc.E14-05-0967.

Cavalli, V., F. Vilbois, M. Corti, M.J. Marcote, K. Tamura, M. Karin, S. Arkinstall, and J. Gruenberg. 2001. The stress-induced MAP kinase p38 regulates endocytic trafficking via the GDI:Rab5 complex. Molecular Cell. 7:421–432. doi:10.1016/S1097-2765(01)00189-7.

Chakraborty, S., M.M. Joseph, S. Varughese, S. Ghosh, K.K. Maiti, A. Samanta, and A. Ajayaghosh. 2020. A new pentacyclic pyrylium fluorescent probe that responds to pH imbalance during apoptosis. Chem. Sci. 11:12695–12700. doi:10.1039/D0SC02623A.

De Chastellier, C., T. Lang, and L. Thilo. 1995. Phagocytic processing of the macrophage endoparasite, Mycobacterium avium, in comparison to phagosomes which contain Bacillus subtilis or latex beads. European Journal of Cell Biology.

de Chastellier, C., and L. Thilo. 1997. Phagosome maturation and fusion with lysosomes in relation to surface property and size of the phagocytic particle. European journal of cell biology. 74:49– 62.

Chen, C., J. Ke, X. Edward Zhou, W. Yi, J.S. Brunzelle, J. Li, E.L. Yong, H.E. Xu, and K. Melcher. 2013. Structural basis for molecular recognition of folic acid by folate receptors. Nature. 500:486–489. doi:10.1038/nature12327.

Coller, S.P., and D.M. Paulnock. 2001. Signaling pathways initiated in macrophages after engagement of type A scavenger receptors. Journal of Leukocyte Biology. 70:142–148. doi:10.1189/jlb.70.1.142.

Dill, B.D., M. Gierlinski, A. Härtlova, A.G. Arandilla, M. Guo, R.G. Clarke, and M. Trost. 2015. Quantitative proteome analysis of temporally resolved phagosomes following uptake via key phagocytic receptors. Molecular and Cellular Proteomics. doi:10.1074/mcp.M114.044594.

Fountain, A., S. Inpanathan, P. Alves, M.B. Verdawala, and R.J. Botelho. 2021. Phagosome maturation in macrophages: Eat, digest, adapt, and repeat. Advances in Biological Regulation. doi:10.1016/j.jbior.2021.100832.

Hajam, I.A., P.A. Dar, I. Shahnawaz, J.C. Jaume, and J.H. Lee. 2017. Bacterial flagellin—a potent immunomodulatory agent. Experimental & Molecular Medicine. 49:e373–e373. doi:10.1038/emm.2017.172.

Han, J., J.D. Lee, L. Bibbs, and R.J. Ulevitch. 1994. A MAP kinase targeted by endotoxin and hyperosmolarity in mammalian cells. Science. 265:808–811. doi:10.1126/science.7914033.

Henry, R.M., A.D. Hoppe, N. Joshi, and J.A. Swanson. 2004. The uniformity of phagosome maturation in macrophages . Journal of Cell Biology. 164:185–194. doi:10.1083/jcb.200307080.

Jankowski, A., C.C. Scott, and S. Grinstein. 2002. Determinants of the phagosomal pH in neutrophils. The Journal of biological chemistry. 277:6059–6066. doi:10.1074/jbc.M110059200.

Kim, S., J. Jang, H. Kim, H. Choi, K. Lee, and I.-H. Choi. 2012. The effects of silica nanoparticles in macrophage cells. Immune Netw. 12:296–300. doi:10.4110/in.2012.12.6.296.

Kim, S.H., J. Kim, and R.P. Sharma. 2004. Inhibition of p38 and ERK MAP kinases blocks endotoxin-induced nitric oxide production and differentially modulates cytokine expression. Pharmacological Research. 49:433–439. 10.1016/j.phrs.2003.11.004.

Linares-Alcántara, E., and F. Mendlovic. 2022. Scavenger Receptor A1 Signaling Pathways Affecting Macrophage Functions in Innate and Adaptive Immunity. Immunological Investigations. 51:1725–1755. doi:10.1080/08820139.2021.2020812.

Mkaddem, S. Ben, M. Benhamou, and R.C. Monteiro. 2019. Understanding Fc receptor involvement in inflammatory diseases: From mechanisms to new therapeutic tools. Frontiers in Immunology. 10. doi:10.3389/fimmu.2019.00811.

Nawaz, F.Z., and E.T. Kipreos. 2022. Emerging roles for folate receptor FOLR1 in signaling and cancer. Trends in Endocrinology and Metabolism. 33:159–174. doi:10.1016/j.tem.2021.12.003.

Nguyen, J.A., and R.M. Yates. 2021. Better Together: Current Insights Into Phagosome-Lysosome Fusion. Frontiers in Immunology. doi:10.3389/fimmu.2021.636078.

Nikolic, D., L. Calderon, L. Du, and S.R. Post. 2011. SR-A ligand and M-CSF dynamically regulate SR-A expression and function in primary macrophages via p38 MAPK activation. BMC Immunology. 12. doi:10.1186/1471-2172-12-37.

Oh, Y.K., and J.A. Swanson. 1996. Different fates of phagocytosed particles after delivery into macrophage lysosomes. Journal of Cell Biology. doi:10.1083/jcb.132.4.585.

Pauwels, A.-M., M. Trost, R. Beyaert, and E. Hoffmann. 2017. Patterns, Receptors, and Signals: Regulation of Phagosome Maturation. Trends in immunology. 38:407–422. doi:10.1016/j.it.2017.03.006.

Rejman, J., V. Oberle, I.S. Zuhorn, and D. Hoekstra. 2004. Size-dependent internalization of particles via the pathways of clathrin- and caveolae-mediated endocytosis. The Biochemical journal. 377:159–169. doi:10.1042/BJ20031253.

Rose, D.M., B.W. Winston, E.D. Chan, D.W. Riches, P. Gerwins, G.L. Johnson, and P.M. Henson. 1997. Fc gamma receptor cross-linking activates p42, p38, and JNK/SAPK mitogen-activated protein kinases in murine macrophages: role for p42MAPK in Fc gamma receptor-stimulated TNF-alpha synthesis. The Journal of Immunology. 158:3433–3438. doi:10.4049/jimmunol.158.7.3433.

Sharma, P., D. Sen, V. Neelakantan, V. Shankar, and S. Jhunjhunwala. 2019. Disparate effects of PEG or albumin based surface modification on the uptake of nano- and micro-particles. Biomaterials science. 7:1411–1421. doi:10.1039/c8bm01545g.

Sharma, P., A. Vijaykumar, J.V. Raghavan, S.R. Rananaware, A. Alakesh, J. Bodele, J.U. Rehman, S. Shukla, V. Wagde, S. Nadig, S. Chakrabarti, S.S. Visweswariah, D. Nandi, B. Gopal, and S. Jhunjhunwala. 2022. Particle uptake driven phagocytosis in macrophages and neutrophils enhances bacterial clearance. Journal of Controlled Release. doi:10.1016/j.jconrel.2022.01.030.

Simeone, R., F. Sayes, E. Lawarée, and R. Brosch. 2021. Breaching the phagosome, the case of the tuberculosis agent. Cellular Microbiology. 23:e13344. 10.1111/cmi.13344.

Thekkan, S., M.S. Jani, C. Cui, K. Dan, G. Zhou, L. Becker, and Y. Krishnan. 2019. A DNA-based fluorescent reporter maps HOCl production in the maturing phagosome. Nature Chemical Biology. 15:1165–1172. doi:10.1038/s41589-018-0176-3.

Vergne, I., J. Chua, and V. Deretic. 2003. Mycobacterium tuberculosis Phagosome Maturation Arrest: Selective Targeting of PI3P-Dependent Membrane Trafficking. Traffic. 4:600–606. 10.1034/j.1600-0854.2003.00120.x.

Westman, J., and S. Grinstein. 2021. Determinants of Phagosomal pH During Host-Pathogen Interactions. Frontiers in Cell and Developmental Biology. doi:10.3389/fcell.2020.624958.

Yates, R.M., A. Hermetter, and D.G. Russell. 2005. The kinetics of phagosome maturation as a function of phagosome/lysosome fusion and acquisition of hydrolytic activity. Traffic. doi:10.1111/j.1600-0854.2005.00284.x.

Yates, R.M., A. Hermetter, G.A. Taylor, and D.G. Russell. 2007. Macrophage activation downregulates the degradative capacity of the phagosome. Traffic. doi:10.1111/j.1600-0854.2006.00528.x.

Yates, R.M., and D.G. Russell. 2005. Phagosome maturation proceeds independently of stimulation of toll-like receptors 2 and 4. Immunity. 23:409–417. doi:10.1016/j.immuni.2005.09.007.

