## Supplementary for "Phagosome Maturation in Macrophages is Enhanced by p38α MAPK Signaling"

This document includes Supplementary Figures S1 – S4

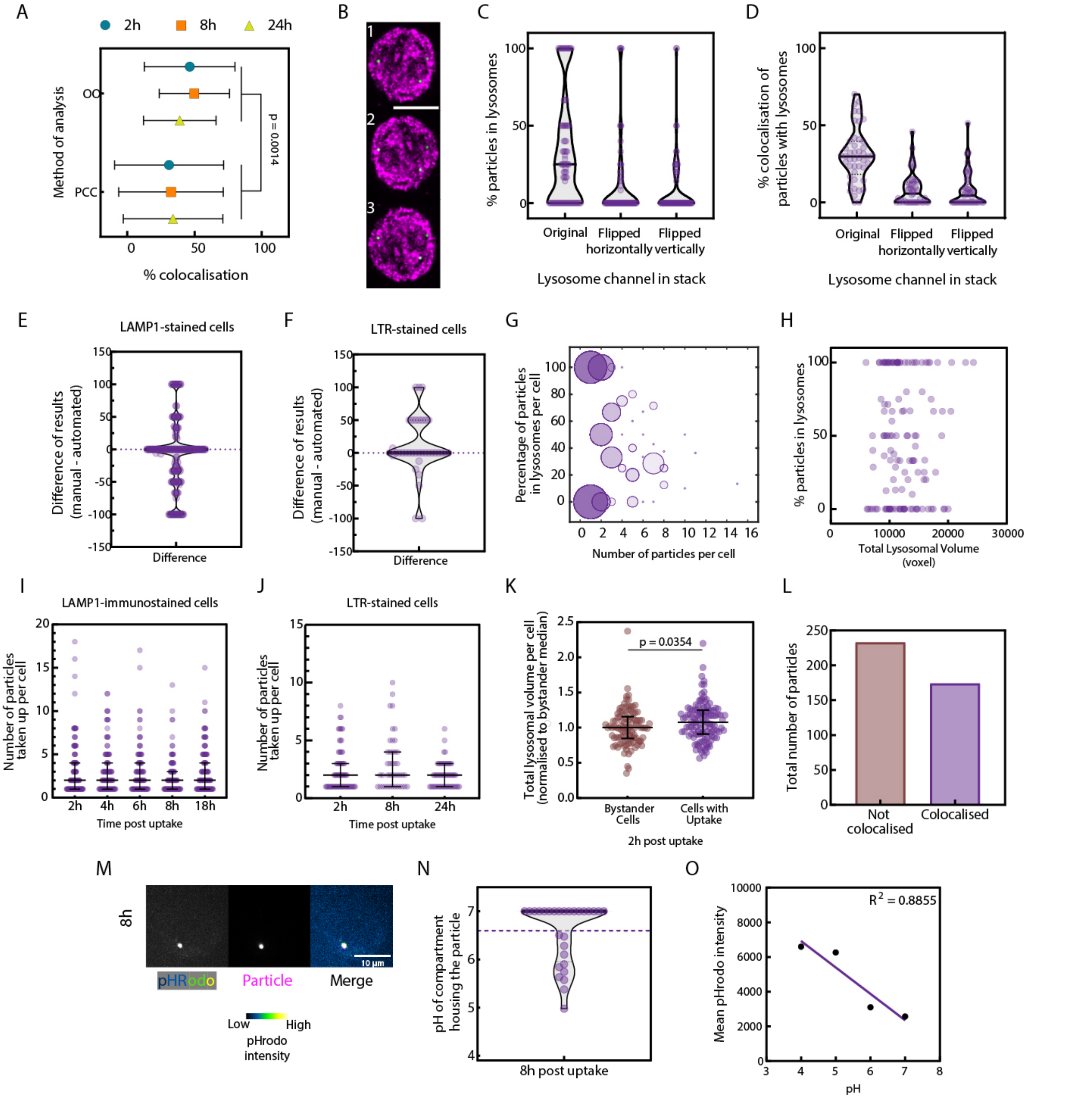


**Supplementary Figure S1: Analysis methods for determining colocalization of particles with lysosomes.** **A –** Comparison of results from the object overlap-based method (OO) and Pearson's correlation coefficient-based method (PCC) in RAW 264.7 cells stained with LTR post internalization of 500 nm sized polystyrene particles. Mean and standard deviation of data are plotted. Ordinary two-way ANOVA, N = 3 independent experiments, 55 cells per group on average. **B –** Representative MIPs of RAW 264.7 cells immuno-stained for LAMP1 (Magenta) 2 hours after uptake of 500 nm sized particles (Green) with the lysosome channel as originally acquired (1), flipped horizontally (2) or flipped vertically (3). Scale bar: 10 µm. **C –** Percentage particles colocalized with LAMP1 signal per cell as quantified using the PCC method in RAW 264.7 cells 2 hours after uptake of particles with the lysosome channel in the acquired image stack kept as original or flipped either horizontally or vertically. n = 80 cells chosen randomly across 3 experiments. **D –** Percentage colocalization of particles with LAMP1 signal per cell as quantified using the OO method in RAW 264.7 cells with the lysosome channel in the acquired image stack kept as the original or flipped either horizontally or vertically. n = 45 cells chosen randomly across 3 independent experiments. For C & D – data are represented on violin plots, where a dark line indicates median and dotted lines indicate quartiles of the data. **E –** Difference in results obtained from either method as indicated plotted for cells randomly sampled across different time points after uptake over

3 independent experiments. n = 199 cells. Cells were immuno-stained for LAMP1 as a lysosome marker. **F –** Difference in results obtained from either method as indicated plotted for cells randomly sampled across different time points after uptake over 2 independent experiments. n = 36 cells. Cells were stained with LTR as a lysosome marker. For E and F – data are represented on violin plots, with dark line indicating median, colored dotted line in the background indicating mean, and dotted black lines indicating quartiles of the data. **G –**No relation observed between number of particles taken up per cell and % of those particles localizing to the lysosome. Data represented on a bubble-plot with the size of a dot indicative of the number of data points at the coordinate. **H –** No relation observed between total lysosome volume in the cell and % internalized particles localizing to the lysosome. Data represented on an XY scatterplot with each dot indicating data from a single cell. For I and J, data are shown for the 18h time-point. 128 cells analyzed across N = 3 independent experiments. **I –**There is no significant difference in the number of particles taken up per cell across different time points after uptake of particles, Kruskal-Wallis Test. N ≥ 3 independent experiments; 116 cells per group on average. Cells were immunostained for LAMP1 **J –** There is no significant difference in the number of particles taken up per cell across different chase durations, Kruskal-Wallis Test. N = 3 independent experiments; 55 cells per group on average. Cells were stained with LTR. For **I** & J – wide bar indicates median, and error bars indicate the inter-quartile range of the data. **K –** Normalized (to bystander median) lysosomal volumes of bystander cells and cells that have taken up particles 2h post introduction of particles. Mann-Whitney test, N = 4 independent experiments, n = 108 cells on average per group. Wide line indicates median and the error bars indicate the inter-quartile range of the data. **L –** Total number of particles pooled across all analyzed cells identified as either localized in the lysosome or not, 2h after uptake. N = 4 independent experiments. **M –** Representative images of RAW 264.7 cells 8 hours after uptake of pHrodo-conjugated 500 nm-sized polystyrene particles. Green Fire Blue LUT – pHrodo, Magenta – fluorescent polystyrene particles. Scale bar – 10 µm. **N –** pH of 500 nm-sized pHrodo-conjugated particles 8h after uptake in RAW 264.7 macrophages. Data represented on a violin plot with the black line indicating median, colored dotted line in the background indicating mean and black dotted lines indicating quartiles of the data. **O –** Standard curve of mean pHrodo intensity with pH, used for quantification of pH in sub-figure **I**. $R^2$ for the linear regression = 0.8855; N = 3 independent experiments.

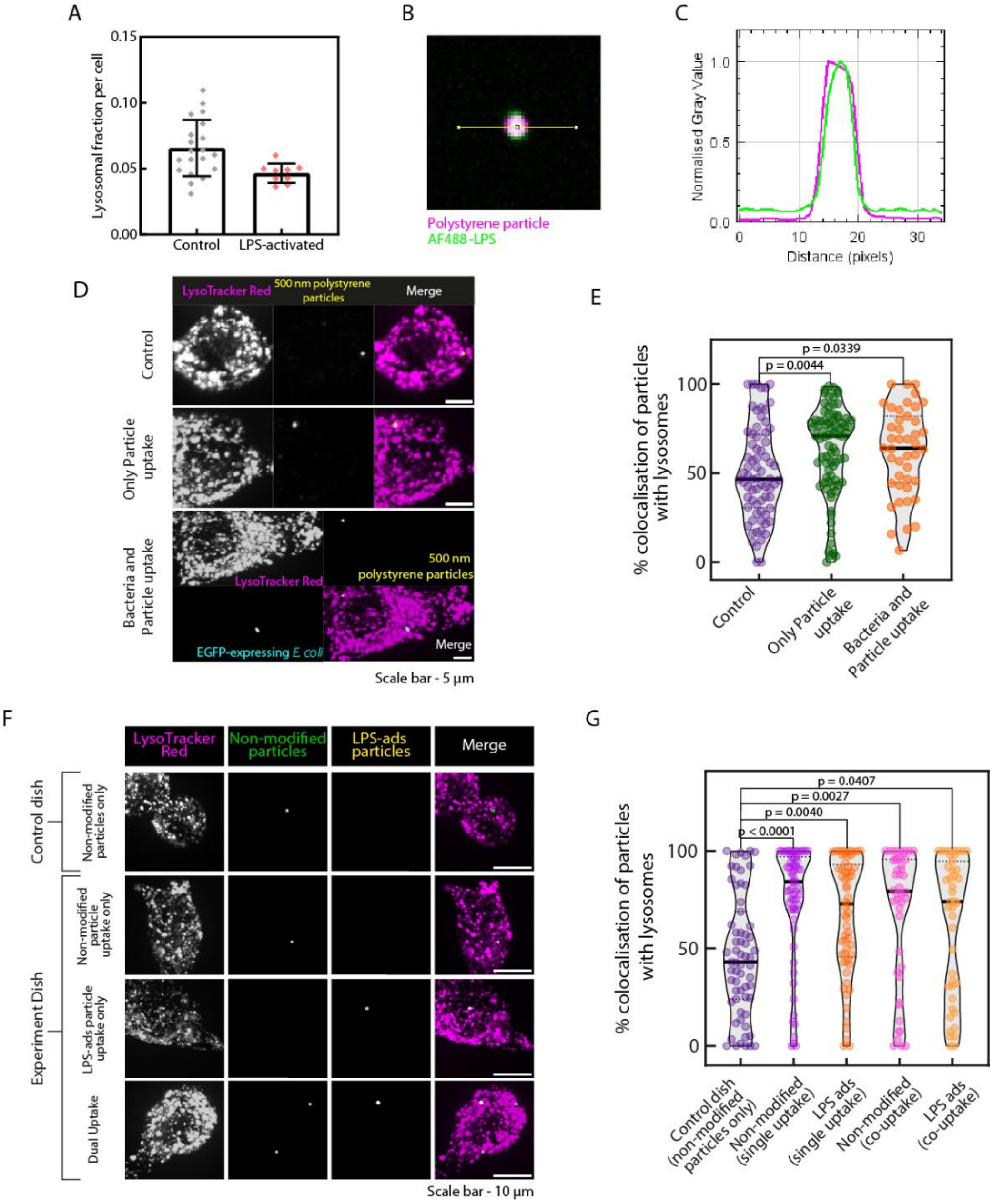


**Supplementary Figure S2: LPS induced signaling enhances phagosome maturation. A –** Fraction of cell volume occupied by lysosomes in control or LPS-activated RAW 264.7 cells. Bar plots show arithmetic-mean with the error bars representing standard deviation of the data. **B –** Representative image of a 500 nm-sized polystyrene particle adsorbed with AlexaFluor488 (AF 488)-LPS and imaged 8h after introduction to cells. Magenta – particle, Green – AF 488 LPS **C –** Normalized intensity profiles of AF 488 and fluorescent particle across ROI drawn in B. Magenta – particles, Green – AF 488 LPS **D –** Representative MIPs of RAW 264.7 macrophages post phagocytosis of either particles alone (Control) or both particles and E. coli, resulting in cells which either took up only particles or both bacteria and particles. (Cells which took up only bacteria are not shown) Magenta – LTR, Yellow – 500 nm particles, Cyan – EGFP-expressing E. coli. Scale bar – 5 µm. **E –** Percentage colocalization of particles with LTR signal per cell 8h post introduction of particles or particles and bacteria as represented in D was calculated. Kruskal-Wallis test with Dunn's multiple comparison post-hoc test. N = 4 independent experiments, n = 65 cells per group on average. **F –** Representative MIPs of RAW 264.7 cells post phagocytosis of only non-modified particles (Control dish) or both non-modified and LPS-adsorbed particles simultaneously (Experiment dish). Magenta – LysoTracker Red, Green – non-modified particles, Yellow – LPS-adsorbed particles; Scale bars – 10 µm. **G –** Percentage of colocalization between particles internalized per cell with LTR signal 8 hours post uptake. Kruskal-Wallis Test with Dunn's multiple comparisons test. N = 3 independent experiments, n = 55 cells per group on average.

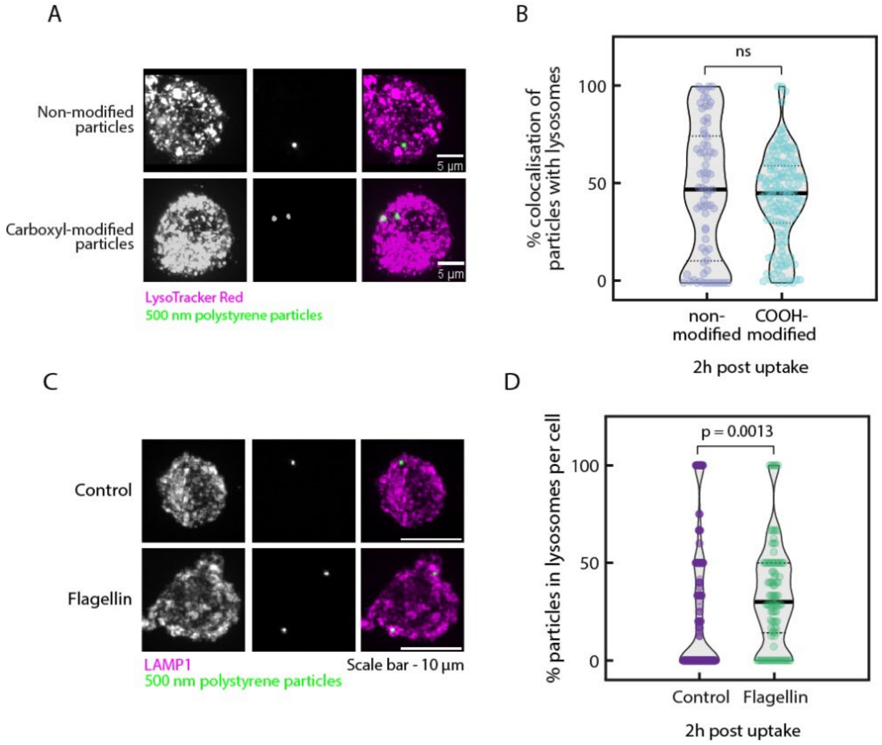


**Supplementary Figure S3: Extents of lysosomal localization of particles with carboxyl surface functional groups or under flagellin-induced signaling. A –** Representative MIPs of RAW 264.7 cells 2h after uptake of non-modified (NM) or -COOH – modified polystyrene particles of 500 nm size. Magenta – LysoTracker Red, Green – 500 nm polystyrene particles. **B –** Percentage colocalization of particles with LTR signal. Mann-Whitney test; ns denotes no significant difference. N = 3 independent experiments, n = 97 cells per group on average. Data are plotted as violin plots with black lines indicating median and dashed lines indicating quartiles of the data. **C –** Representative MIPs of either control cells or cells treated with Flagellin for 18h before uptake of polystyrene particles. Magenta – LAMP1, Green – particles, Scale bar – 10 µm. **D –** Percentage of particles taken up per cell that colocalize with LAMP1 2h after uptake as represented in C. Mann-Whitney test, N = 3 independent experiments, n = 191 cells per group on average. Data are plotted as violin plots with black lines indicating median, dashed lines indicating quartiles, and colored lines indicating arithmetic-mean of the data.

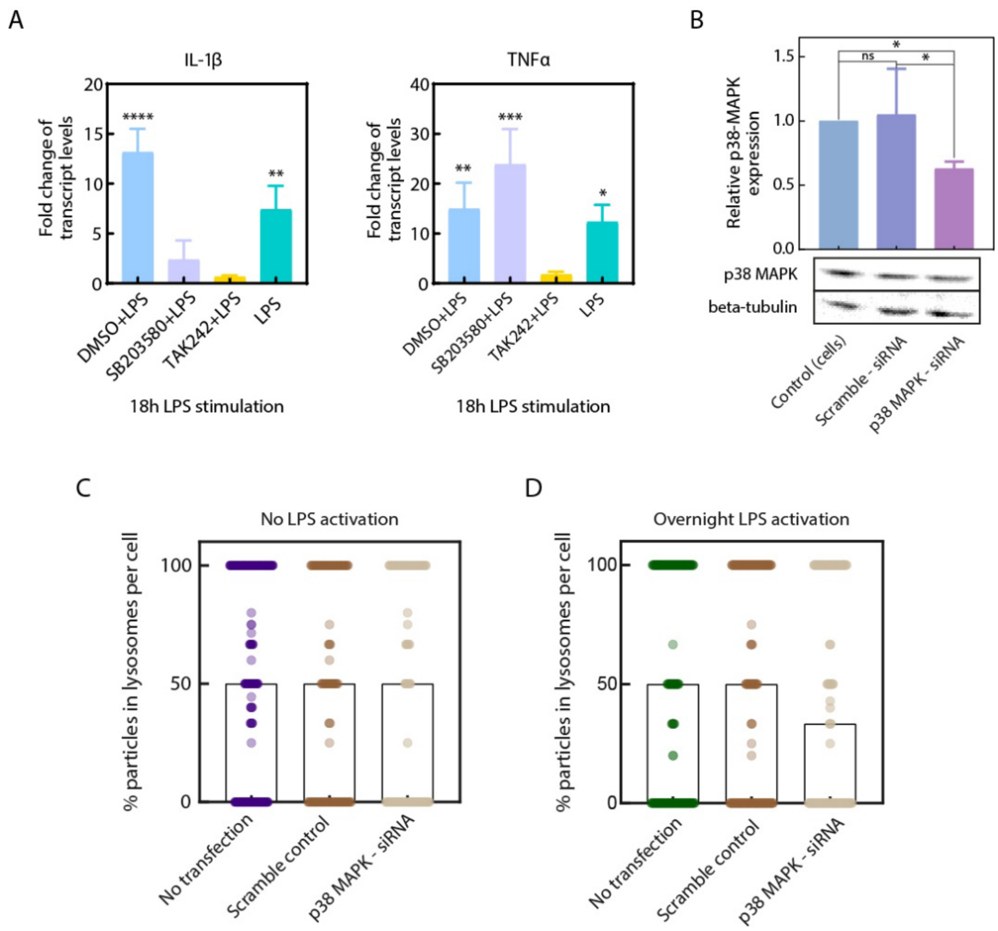


**Supplementary Figure S4: Inhibition of p38 MAPK activity abrogates LPS-induced enhancement of phagosome maturation. A –** Relative transcript levels compared to untreated, control cells (normalized to Gapdh transcript levels) of cytokines IL-1β and TNFα in RAW 264.7 cells under indicated treatments – either 18 hours of 100 ng/mL treatment or the same preceded by 10 μM SB203580 or 1 μM TAK242 or equivalent DMSO treatment for 1 hour. **B –** Quantification of Western blot assay along with a representative blot image to verify the siRNA-mediated knockdown of p38 MAPK in RAW264.7 cells. N = 5 independent experiments, One-way ANOVA with Tukey's post-hoc test, ns = no significant difference. For A, B – bar plots indicate mean of the groups with error bars indicating standard deviation. **** - $p < 0.0001$, *** - $p < 0.001$, ** - $p < 0.01$, * - $p < 0.05$. **C –** Percentage of particles taken up per cell that localize to lysosomes, 2h post uptake in RAW264.7 cells which were transfected with scramble-siRNA or siRNA against p38 MAPK, or not subjected to transfection as a control. N = 3 independent experiments, n = 169 cells on average per group. **D –** Percentage of particles taken up per cell that localize to lysosomes, 2h post uptake in RAW 264.7 cells which were treated overnight with 100 ng/mL LPS 48 hours after transfection with either scramble-siRNA or siRNA against p38 MAPK, or not subjected to transfection as a control. N = 3 independent experiments, n = 159 cells on average per group. For C, D – data are represented on bar plots with the bars indicating medians of the groups.
